# Murine models for triple-negative breast cancer with differential responsiveness to immunotherapy

**DOI:** 10.1101/2025.09.18.677171

**Authors:** Thomas J Kalantzakos, Yufan Zhou, Xingyu Liu, Joshua Proehl, Cameron Durfee, Ian Tamayo, Nuri Alpay Temiz, Benjamin Troness, Anusha Soni, Harshita B Gupta, Reuben S Harris

## Abstract

Breast cancer is the most common cancer diagnosis in women. Clinical studies with triple-negative breast cancer (TNBC) are encouraging for immunotherapy combined with chemotherapy (anti-PD-1 with paclitaxel and/or carboplatin). However, additional clinical advances may be pursued more rapidly with assistance from preclinical TNBC models including syngeneic mammary tumor cell lines. Here, we report two mammary tumor cell lines that exhibit differential responsiveness to immunotherapy *in vivo*. Spontaneous mammary tumors from C57BL/6J MMTV-Cre Trp53^fl/+^ animals were passaged serially in cell culture and *in vivo* in the mammary fat pad of fully wildtype animals. The resulting lines, MM001i and MM008i, lost *Trp53* and formed 1000 mm^3^ tumors in the mammary fat pad within 21-28 days. Despite originating from the same genetic background, these lines exhibit differential responses to immunotherapy. For anti-PD-1 therapy, MM001i is poorly responsive and MM008i is strongly responsive with near-complete tumor regression. In comparison, both MM001i and MM008i respond rapidly to anti-CTLA-4 therapy. Both models express unique tumor antigens as evidenced by immunity to subsequent engraftments. Primary MM008i tumors exhibit greater T cell infiltration, and CD8-positive T lymphocytes are required for anti-PD-1 responses. These TNBC models are promising for further mechanistic studies and testing future single and combinatorial therapies.

## Introduction

Breast cancer is the most commonly diagnosed malignancy worldwide and a leading cause of cancer-related death in women.^1, 2^ Clinically, breast cancer is characterized based on expression of estrogen receptor (ER), progesterone receptor (PR), and/or human epidermal growth factor receptor 2 (HER2). Triple negative breast cancer (TNBC) lacks expression of these three markers, accounts for 15-20% of breast cancer diagnoses, and is associated with poor clinical outcomes.^3^ For example, 30-40% of TNBC patients exhibit tumor recurrence within 3-5 years of the initial diagnosis and treatment.^4, 5^ The most common genetic defect in TNBC is mutation of *TP53* (hereafter *p53*) in 75-80% of cases,^6, 7^ with most tumors incurring full loss-of-function alleles.^8^ TNBC tumors are highly heterogeneous and often subdivided into six molecular clusters: basal-like 1, basal-like 2, immunomodulatory, mesenchymal, mesenchymal stem-like, and luminal androgen receptor subtypes.^9^

Several clinical trials have demonstrated the benefit of immunotherapy approaches in TNBC, particularly through monoclonal antibody (mAb) targeting of programmed cell death protein 1 (PD-1) and cytotoxic T-lymphocyte antigen 4 (CTLA-4). The efficacy of immunotherapy, particularly anti-PD-1 antibody therapy (αPD-1), in combination with chemotherapy, is becoming established in both primary and metastatic TNBC settings. In the KEYNOTE-522 clinical trial, adding αPD-1 to neoadjuvant chemotherapy (paclitaxel and carboplatin followed by doxorubicin/epirubicin and cyclophosphamide) in early TNBC improved pathological complete response from 51% to 65%.^10^ Further, the KEYNOTE-355 trial, conducted on advanced TNBC patients, demonstrated a robust increase in overall survival when combining αPD-1 and chemotherapy in patients with high programmed cell death-ligand 1 (PD-L1) expression.^11^ Emerging studies are also evaluating combination immunotherapy approaches. For example, the BELLINI trial is investigating dual inhibition of PD-1 and CTLA-4, showing signs of efficacy in early-stage TNBC.^12^ Additionally, bi-specific antibodies targeting PD-1 and CTLA-4 combined with paclitaxel have shown survival benefits for TNBC patients in the metastatic setting.^13^ Additional trials are ongoing that combine immunotherapies with radiation therapy and/or chemotherapy (*e.g*., NCT04683679, NCT05491226, NCT06472583). However, despite these recent advances, a large subset of TNBC patients still suffers from disease progression, which warrants further mechanistic and therapeutic studies.

Syngeneic murine models are useful for preclinical evaluation of candidate therapeutics *in vivo.* To date, only a few murine syngeneic models for TNBC have been reported. The first, E0771, was derived in 1955 from a spontaneously occurring breast adenocarcinoma in a C57BL/6 background.^14^ Treatment of this model with αPD-1 leads to partial tumor regression^15^ and enhanced efficacy when combined with other agents.^16^ αPD-1 and αCTLA-4 therapies also associate with increases in circulating CD8^+^ T lymphocytes in this model.^16, 17^ However, it is an older model and reports conflict on several key properties including genotype and hormone receptor status.^18–20^ Another commonly used line is 4T1, isolated from a metastatic tumor in a BALB/c animal.^21^ However, this line is poorly responsive to immunotherapy due at least in part to a Th2-biased immune response.^22, 23^ Other models, such as PyMT,^24^ AT-3,^25^ and WT145^26^ are additionally complicated by viral transgenes and/or chemical exposures.

Here, we report the development of two new cell lines, MM001-immunoconditioned (MM001i) and MM008i, derived from mammary tumors that arose on C57BL/6J background. Briefly, the primary mammary tissues were engineered to be heterozygous for *p53*, but the second copy was lost during tumor development in both cell lines. Serial passage in culture and *in vivo* resulted in MM001i and MM008i, which form 1000 mm^3^ tumors approximately 21-28 days after engraftment. Both lines are negative for ER, PR, and HER2 (triple-negative) and provoke anti-tumor immune responses, as evidenced by immune memory formation and acquired resistance to additional challenge by the same tumor cell line. Both lines also respond to αCTLA-4 immunotherapy, but only MM008i responds to αPD-1 immunotherapy. This differential responsiveness is dependent on CD8^+^ T lymphocytes and may be attributable to improved immune cell infiltration in MM008i tumors (CD45^+^ immune cells, CD8^+^ T lymphocytes). Taken together, these syngeneic tumor lines constitute new models for TNBC and provide a foundation for additional studies on molecular mechanisms and responsiveness of different immunotherapies.

## Materials and Methods

### Mouse care

All animal studies were approved by the University of Texas Health San Antonio (UT Health) Institutional Animal Care and Use Committee (IACUC Protocol ID: 20220056AR) and conducted in accordance with ARRIVE guidelines 2.0.^27^ The study period was 2023-2025. “Guide for the care and use of laboratory animals” guidelines were followed, and the study adhered to the principles of 3Rs (Replace, Reduce, and Refine). The Lab Animal Resources (LAR) facilities at UT Health are AAALAC (accredited unit: #000341) and USDA (registration number: 74-R-0071) approved. 24/7/365 LAR staff include veterinarians, facility staff, and animal technicians for services including complete husbandry, routine monitoring, examination of mice health and supportive care. 3-5 animals per cage were housed in specific pathogen-free conditions maintained at 22°C under a standard 12-hour light/dark cycle and given food and water *ad libitum*. Initial breeder pairs for WT (Jax strain 000664) and NOD.Cg-Prkdc^scid^Il2rg^tm1Wj1^/SzJ [nonobese diabetic/severe combined immunodeficiency (NOD/SCID)/IL2Rγ KO, NSG, Jax strain 005557] mice were purchased from Jackson Laboratory (Bar Harbor, ME, USA) and then propagated in-house. NSG mice were maintained under sterile conditions. 8-10 weeks-old mice were enrolled for tumor studies. All immunocompetent mice were on C57BL/6J background, and only females were used for mammary tumor experiments.

Mice were checked one day post-injection, and then twice a week for tumor growth, morbidity, weight loss, and dehydration. Mice were euthanized if they were in distress, lost >20% body weight, showed a body score <2, and/or reached experimental end points such as 1000 mm^3^ tumor volumes. The signs of distress included but were not limited to dehydration, cachexia, weakness, ruffled body fur, or hunched back. Euthanasia was done as recommended by the American Veterinary Medical Association guidelines for the Euthanasia of Animals. Briefly, mice were subjected to inhalation of 99% pure CO_2_ with a fill rate of 30-70% chamber volume/minute for 2 minutes followed by cervical dislocation. Anesthesia was not used for the experiments performed for this study.

### Tumor harvesting from mice

*MMTV-Cre p53*^fl/+^ mice were bred initially at the University of Minnesota and subsequently at UT Health prior to the present studies. Genotypes were confirmed via PCR^28^ using primers: 1F: 5’-TAACACTCTTGGGATGCTGAG-3’, 10F: 5’-AAGGGGTATGAGGGACAAGG-3’, and 10R: 5’-GAAGACAGAAAAGGGGAGGG-3’. Spontaneously arising primary mammary tumors in these mice were extracted under sterile conditions, and single-cell suspensions were made by tumor dissection and mechanical dissociation. Cells were transferred to a cell culture flask in RPMI-1640 (Thermo Fisher Scientific, Waltham, MA, USA) supplemented with 10% fetal bovine serum (FBS, Thermo Fisher Scientific) and 1% penicillin-streptomycin (complete medium, Thermo Fisher Scientific), under standard culturing conditions (37°C, 5% CO_2_). Once the cells reached 80% confluence, cells were harvested by 0.05% trypsin-EDTA (Gibco, Thermo Fisher Scientific) for engraftment into mice. Freeze downs were made simultaneously in 90% FBS/10% DMSO (Sigma-Aldrich, St. Louis, MO, USA). All cell lines were confirmed to be free of mycoplasma contamination by detection kit (catalog no. LT07-318, Lonza Bioscience, Walkersville, MD, USA) prior to freezing or experimental use. Cultured cells in passage 3 or 4 were used for subsequent characterization and engraftment studies, which we have found yield reproducible results across multiple biological replicates.

### Serial passage to develop conditioned cell lines

Cells were trypsinized, washed with endotoxin-free, sterile phosphate-buffered saline (PBS) and combined at 1:1 ratio with Cultrex Basement Membrane Extract, PathClear (R&D Systems, Minneapolis, MN, USA) for a final cell count as indicated (2.5 or 5 x 10^6^ cells) in a final volume of 100 μL per injection. Cells were injected subcutaneously or into the 4^th^ inguinal mammary fat pads of mice. Initial engraftments were conducted in both WT and NSG mice. As tumors successfully engrafted in both strains, subsequent engraftments for serial passage were conducted in WT mice alone. At day 7, tumors were measured using Vernier caliper (Thermo Fisher Scientific) and then monitored twice a week until reaching a volume of 1000 mm^3^ using the equation V=0.5 x L x W^2.29^ Tumors reaching 1000 mm^3^ were subsequently isolated post-humane mouse euthanasia. The process was repeated for at least two rounds per cell line.

### Flow cytometry

MM001i or MM008i cells were plated overnight in 6-well plates in complete medium (3 x 10^5^ cells/well). Cells were then incubated with fresh complete medium supplemented with either vehicle or interferon-gamma (IFN-γ) (R&D Systems) for 24 hours. Cells were harvested and counted using the Countess cell counter (Thermo Fisher Scientific). 1 x 10^6^ cells per condition were analyzed for subsequent steps. Cells were washed with PBS, stained for live/dead dye (Zombie Yellow Fixable Viability dye) in PBS for 40 min at 4°C in the dark. Cells were washed in 1x PBS and incubated with anti-CD16/32 (BioLegend, San Diego, CA, USA) in flow buffer (PBS + 2% FBS) to prevent non-specific antibody binding for 30 minutes at 4°C, followed by incubation with anti-mouse PD-L1 (PE, clone 10F.9G2), H-2KB (Pacific Blue, clone AF6-88.5), or matched isotype controls (BioLegend). Cells were fixed in 0.4% buffered formalin. Cells were acquired using BD LSRFortessa (BD Biosciences, San Juan, CA, USA) and analyzed using FACSDiva and FlowJo 10.10.0 (BD Biosciences). Positivity thresholds were determined based on isotype control staining.

### Immunotherapy experiments

2.5 x 10^6^ MM001i or MM008i cells were suspended in 50 μL endotoxin-free PBS. An equal volume of basement membrane was added to reach a final volume of 100 μL/injection. For all immunotherapy experiments, mice were randomized manually to equalize numbers across treatment groups. Mice with no palpable tumor by day 7 (tumor vol < 200 mm^3^) were excluded due to likely failed injection (<25% of injections). Each mouse received two injections (one per inguinal mammary fat pad), and the sample size for each group was based on the total number of animals enrolled for each condition. On day 7, an initial tumor measurement was taken as previously described. Tumors were monitored twice a week, and the mice were euthanized when tumors reached 1000 mm^3^, or when they displayed signs of distress. 7 days post-injection, each mouse received an intraperitoneal (i.p.) injection of 100 μg of αPD-1 (catalog no. BE0146, clone RMP1-14, BioXCell NH, USA), αCTLA-4 (catalog no. BE0131, clone 9H10, BioXCell), a combination of the two, or their respective isotype controls (catalog # BE0089 and BE0087, BioXCell). Mice injected with MM001i received two additional treatments on days 12 and 19.

### Memory response experiments

WT mice with complete tumor regression in response to immunotherapy were enrolled in memory rechallenge experiments. For MM001i, this consisted of one mouse previously treated with αPD-1 and four that received αCTLA-4 or a combination of αPD-1 and αCTLA-4. Additionally, five animals were enrolled that completely cleared MM008i tumors from each treatment group. 90 days after the last tumor evaluation, mice were reinjected with 2.5 x 10^6^ cells of the initial cell line at the same inguinal mammary fat pad. No further treatments were administered. Tumors were measured beginning 7 days post-injection and monitored as above until the endpoint was reached. The control group consisted of age-matched, naïve, previously unchallenged WT mice injected with the same tumor cell line as the rechallenge group but not treated with any immunotherapy agent.

### Immunohistochemistry

IHC was performed as previously described.^30^ In brief, FFPE tissues and cell pellets were sectioned at 4 μm and mounted on positively charged glass slides. The slides were deparaffinized at 65°C for 20 minutes, rehydrated with three 5-minute washes in CitriSolv (Decon Labs, Swedeland, Pennsylvania, USA) followed by 3-minute washes in graded ethanol to 80%, and finished with a 5-minute wash in running water. Antigen retrieval was performed in 1x Reveal Decloaker (Biocare Medical, Pacheco, California, USA) by steaming for 35 minutes and returned to room temperature with 30 minutes off the steamer. The slides were washed for 5 minutes in running water and then in tris-buffered saline with 0.1% tween (TBST, Thermo Fisher Scientific). Endogenous peroxidase activity was suppressed by soaking slides in 3% H_2_O_2_ diluted in TBST for 10 minutes, followed by a 5-minute wash in running water. The slides were soaked in Background Sniper (Biocare Medical) for 15 minutes, immediately followed by overnight incubation of primary antibody diluted using 10% Sniper in TBST at 4°C. The following primary antibodies were used: αER (Catalog no. AB32063, 1:2000, Abcam, Cambridge, UK), αPR (Catalog no. 25871-AP-1, 1:10000, ProteinTech, Rosemont, Illinois, USA), αHER2 (Catalog no. CST2165S, 1:1000, Cell Signaling Technologies, Danvers, Massachusetts, USA), αCD3 (Catalog no. AB135372, 1:250, Abcam), αCD8 (Catalog no. AB209775, 1:600, Abcam) and αCD19 (Catalog no. AB317335, 1:600, Abcam). After overnight incubation, the slides were brought to room temperature and washed in TBST for 5 minutes. Next, they were incubated with Novolink Polymer (Lieca Biosystems, Deer Park, Illinois, USA) for 30 minutes and developed using Novolink DAB substrate kit (Lieca Biosystems) for 5 minutes. Slides were immediately washed in running water to quench substrate development and counterstained for 10 minutes using Mayer’s hematoxylin solution (Electron Microscopy Sciences, Hatfield, Pennsylvania, USA). Slides were then dehydrated in degraded alcohols and CitriSolv and then cover-slipped using Permount mounting media (Fisher Scientific).

### Immune cell infiltration analysis by IHC

MM001i and MM008i sections were used to assess the infiltration of T and B lymphocytes in n=4-5 tumors. T cells and B cells were visualized using αCD3 and αCD19 antibodies respectively, as described above. T cell and B cell densities were calculated using QuPath v0.5.1 bioimage analysis software using the positive cell detection algorithm. 5 random uniform regions of interest (ROIs) were selected within each tumor section and cell density was averaged for all ROIs per tumor.

### Immune cell infiltration analysis by flow cytometry

Mice were euthanized, and tumors were harvested once the mean group volume reached 750 mm^3^. Tumors were digested, and single-cell suspensions were made using 0.3 mg Liberase (Thermo Fisher Scientific) and 0.2 mg DNase I (Sigma-Aldrich) per tumor in incomplete RPMI-1640. Tumors were mechanically dissociated, minced and incubated with shaking at 37°C for 30 minutes. Cell suspensions were harvested, passed through 70 μM strainers, and counted. 5 x 10^6^ cells were stained for surface markers as above. Cells were incubated with αCD16/32 to prevent non-specific antibody binding as above, followed by staining with αCD45 (FITC), αCD3 (PE-Cy7), αCD8 (APC-Cy7), αCD4 (Bv786), αCD62L (Bv711), αPD-1 (PerCP-Cy5), αCD44 (Pacific Blue), αTIM3 (APC), or αGr1 (Bv786), αCD86 (APC-Cy7), αCD11c (PE), αCD11b (PerCP-Cy5), αB220 (AF700), αNK1.1 (APC) post live/dead staining (Zombie Yellow Fixable Viability dye). All antibodies and Zombie Yellow Fixable Viability dye were purchased from BioLegend. Cells were washed, fixed in 0.4% buffered formalin, and analyzed as mentioned above.

### CD8^+^ neutralization experiments

2.5 x 10^6^ MM008i cells were injected into the mammary fat pads of immunocompetent WT mice as previously described. On day 6, initial tumor measurements were taken, and mice were randomized into groups. On days 6, 9, and 12, each mouse received an i.p. injection of 200 μg of αCD8 antibody (catalog no. BE0004-1, clone 53-6.7, BioXCell) or its isotype control (catalog no. BE0089, clone 2A3, BioXCell). On day 7, each mouse received a single i.p. injection of 100 μg of αPD-1 or its isotype control, as previously described. Tumors were measured biweekly, and mice were euthanized at day 25 or when tumor volumes reached 1000 mm^3^. At day 13 and before complete tumor regression, a small subset of tumors representing each treatment group were isolated to assess immune infiltration of CD8^+^ T lymphocytes by immunohistochemistry.

### Bulk RNA-sequencing of MM001i and MM008i cell lines and tumors

RNA from two biological replicates each of MM001i and MM008i grown in culture were isolated using the Qiagen RNeasy Mini Kit (Qiagen, Hilden, Germany). Three biological replicates of RNA from MM001i and MM008i tumors each were isolated using the Qiagen AllPrep DNA/RNA Mini kit (Qiagen). Tumor tissues were mechanically homogenized in lysis buffer containing β-mercaptoethanol using a glass Dounce homogenizer, followed by repeated aspiration through a syringe for disruption. RNA quality was confirmed by Nanodrop spectrophotometer. Stranded RNA-seq libraries were prepared in collaboration with the Genome Sequencing Facility at UT Health for paired-end 100 bp sequencing. The sequencing raw reads were mapped to the mm10 *Mus Musculus* reference genome with HISAT2 v2.1.0^31^, and gene counts were calculated with htseq-count. Comprehensive gene expression data for cultured cells and tumors of each cell line are available in Supplementary Table 1. Enrichment of KEGG signaling was conducted using the top 1000 expressed genes of each cell line in each condition, with specific focus on immune pathways, cancer cell pathways, and epithelial cell pathways. The input genes and enriched pathways for MM001i and MM008i cultured cells and tumors are available in Supplementary Table 2. Differentially expressed genes in both datasets were identified with DEseq2 v1.40.2^32^ by cutoff of log2 fold change > 2 and adjusted *P* value < 0.01 for tumors and < 0.05 in culture (*P* value by Wald test and *P* value adjustment by Benjamini-Hochberg adjustment).

### Single-cell expression profiling

Three mammary tumors each of MM001i and MM008i were collected from six individual mice. Single-cell suspensions were prepared for the 10X Genomics Chromium system using a tumor dissociation kit (catalog no. 130-096-730, Miltenyi Biotec, San Diego, CA, USA) at a final concentration of 1000 cells/uL. Single-cell RNA sequencing was performed in collaboration with the Genome Sequencing Facility at UT Health according to the technical manual. Raw sequencing reads were aligned to the mouse reference genome mm10 using 10x Genomics Cell Ranger. Feature-barcode matrices were generated with Cell Ranger’s count pipeline. Subsequent analysis was performed in Seurat (v4.4.0)^33^ using the following workflow: a Seurat object was created with CreateSeuratObject using thresholds of min.cells = 3 (genes detected in ≥3 cells) and min.features = 500 (cells expressing ≥500 genes). Cells with >30% mitochondrial transcript content (percent.mt < 30) were excluded via subset. For MM001i, a total of 57,866 cells were profiled and, after filtering, 40,645 cells remained in the analysis. For MM008i, 56,804 cells were profiled and 39,726 cells remained after filtering. For each tumor model, cells were pooled and clusters identified using FindClusters with a resolution parameter of 0.5. Cell types were identified manually with curated cell markers: T cells: *Cd3g*, *Cd3d*, *Cd3e*, *Cd247*; B cells: *Cd19*, *B220*; Macrophages: *Cd74*, *Cd14*, *Csf1r*; Endothelial cells: *Esam*, *Pecam1*; Stromal cells: *Fn1*, *Fap*, *Acta2*, *S100a4*; Basal cells: *Krt14*, *Krt5*, *Krt17*; Epithelial cells: *Cdh1*, *Krt8*, *Krt18*, *Krt19*; Cancer cells: *Acta2*, *Aldh1a3*, *Cd24a*, *Cdh1*, *Cdh2*, *Erbb2*, *Kit*, *Krt7*, *Mfge8*, *Myc*, *Pgr*, *Prlr*, *Tjp1*, *Top2a*, *Twist1*.

### Nomenclature

Throughout the manuscript, gene and transcript names are denoted with italics and protein names with non-italics. All gene/protein names are capitalized fully regardless of species, except for tumor gene expression analyses which use mouse nomenclature (first letter capitalized).

### Statistical analysis

Calculations were done using Microsoft Excel software, and statistical analyses were performed using GraphPad Prism software version 10.3.1 (GraphPad, La Jolla, CA, USA). Data were plotted as means ± standard deviation. For tumor growth, Two-way analysis of variance (ANOVA) was used to compare replicate means between multiple groups. Unpaired Student’s *t*-test was used to compare individual means. *P* < 0.05 was considered statistically significant.

## Results

### Generation of murine mammary tumor cell lines MM001i and MM008i

Two spontaneously arising mammary tumors were harvested from independent C57BL/6J *MMTV-Cre p53*^fl/+^ mice, and cell lines were generated by serial passage in culture and *in vivo* (Supplementary Figure 1A; Materials and Methods). Primary tumors were isolated under sterile conditions and expanded in culture to obtain sufficient cells for cryopreservation and engraftment studies. 5 x 10^6^ cells were injected into the mammary fat pad or subcutaneously in severely immunocompromised NSG mice, and tumor formation occurred rapidly within 14-21 days as expected in most animals regardless of injection site (not shown). However, parallel engraftments of 5 x 10^6^ cells into the same anatomical locations in fully immunocompetent C57BL/6J (WT) animals only led to tumor formation in the mammary fat pad (Supplementary Figure 1). Representative tumors from the first mammary fat pad engraftment were expanded in culture and used in a second mammary fat pad engraftment. After two (MM001i) or three (MM008i) rounds of culturing and engraftment, these cell lines formed stable and reproducible mammary tumors (Supplementary Figure 1D,H). For post-tertiary mammary fat pad engraftments in WT mice, 2.5 x10^6^ cells were sufficient to form tumors in 21-28 days with at least 70% efficiency. Therefore, this cell number was used for all subsequent experiments.

### MM001i and MM008i tumors are potential models for human TNBC

As these tumor cell lines had been generated from mice with mammary-specific heterozygosity for *p53*, PCR was used to determine the genotype of this locus (Supplementary Figure 2A; primers listed in methods).^28^ Although both MM001i and MM008i harbored one *p53* null allele (*i.e*., *loxP*-to-*loxP* recombined) as anticipated, the WT allele could not be detected. Taken together, these PCR results indicated that both MM001i and MM008i are functionally *p53* homozygous null (likely through loss-of-heterozygosity early in tumor development).

Mammary tumor cell lines lacking functional p53 are typically also negative for the hormone receptors, ER, PR, and HER2.^34, 35^ To visualize these cell surface markers, representative tumors from both MM001i and MM008i were formalin-fixed and paraffin-embedded for immunohistochemical staining. MM001i and MM008i are both triple-negative at the protein level, lacking positive nuclear staining of ER^36^ and PR proteins^37^ as well as cell surface staining of HER2^38^ (Supplementary Figure 2B).

Next, we used flow cytometry to quantify the surface expression of pro-tumorigenic, immune co-signaling molecule PD-L1 on our cell lines at a basal levels and following 24 hrs induction with interferon-gamma (IFN-γ)^12, 29^. MM008i showed higher expression of PD-L1 at basal levels as well as after IFN-γ treatment compared to MM001i (Supplementary Figure 2C). However, cell surface major histocompatibility complex-I (H-2KB) expression was similar between both cell lines regardless of IFN-γ treatment (Supplementary Figure 2D).

### MM001i and MM008i respond differentially to αPD-1 but similarly to αCTLA-4

After establishing a robust mammary fat pad engraftment protocol, WT females were split randomly into subgroups and treated with the two most commonly used immunotherapy agents, αPD-1 and αCTLA-4, to gauge therapy responsiveness. After engrafting on day 0 as above, treatment with αPD-1, αCTLA-4, or the combination began on day 7, once tumors reached approximately 200 mm^3^. MM001i engrafted animals also received treatments on days 12 and 17 (Figure 1A). These tumors responded poorly to αPD-1 but regressed completely with αCTLA-4 monotherapy as well as with αPD-1 + αCTLA-4 combination therapy (Figure 1B). Non-responsiveness was observed as expected in severely immunocompromised NSG mice (Figure 1C), indicating that these immunotherapy outcomes are immune response-dependent.

**Figure 1.**
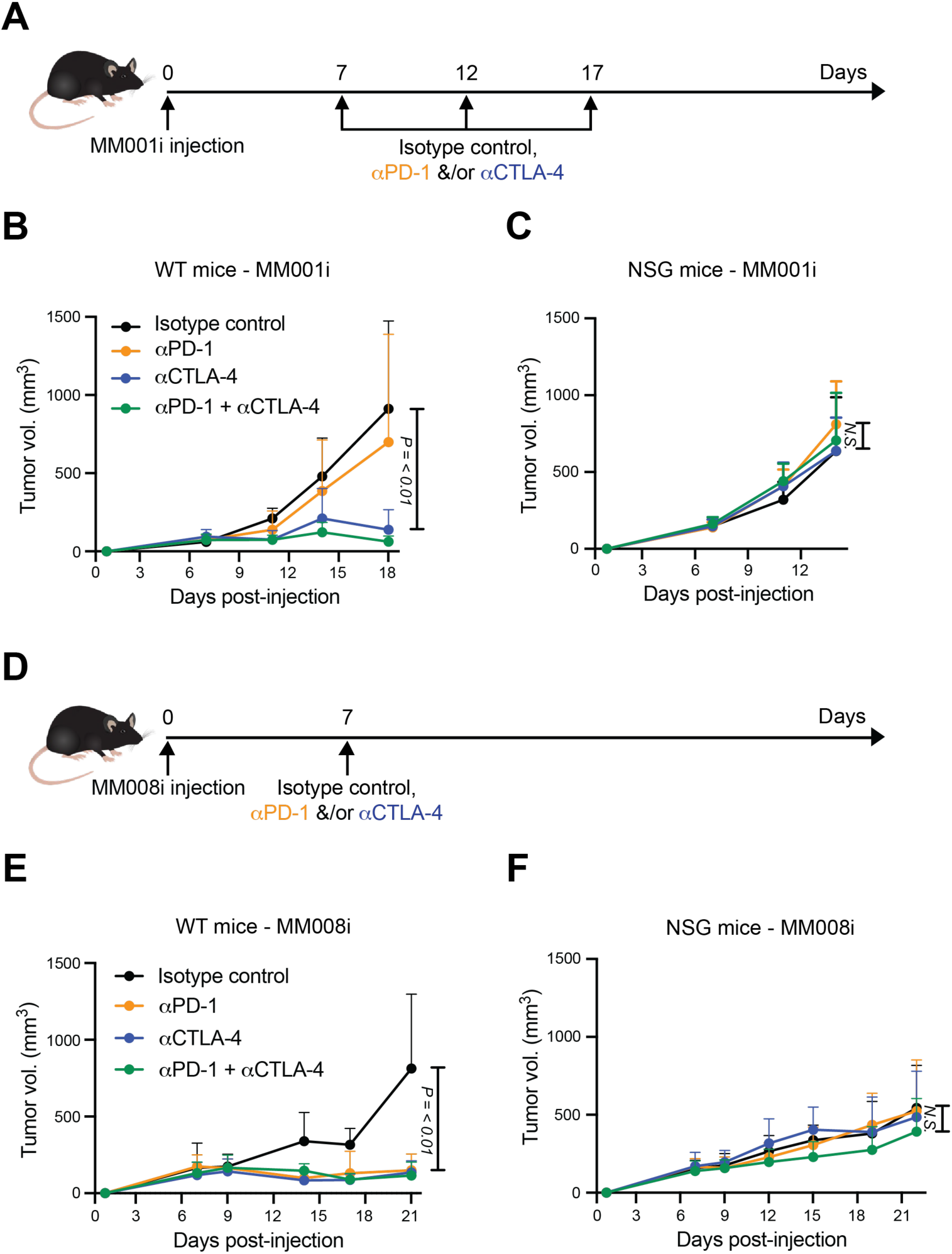
MM001i and MM008i tumors exhibit differential immunotherapy responsiveness. (**A**) Timeline of MM001i tumor injection and treatment with αPD-1 ± αCTLA-4 or isotype controls. (**B-C**) Tumor growth curves of MM001i following injection into the mammary fat pads of WT or severely immunocompromised (NSG) female mice, respectively, in the indicated treatment groups. n=6 (WT) or 4 (NSG) animals per group with two tumors per mouse; *P*-values using two-way ANOVA. (**D**) Timeline of MM008i tumor injection and treatment with αPD-1 ± αCTLA-4 or isotype controls. (**E-F**) Tumor growth curves of MM001i following injection into the mammary fat pads of WT or NSG female mice, respectively, in the indicated treatment groups. n=6 (WT) or 4 (NSG) animals per group with two tumors per mouse; *P*-values using two-way ANOVA.

MM008i tumors were treated with a single dose on day 7 (Figure 1D). Additional doses were not necessary because tumors regressed rapidly following this single treatment. MM008i tumors responded equally well to both αPD-1 and αCTLA-4 monotherapies, ultimately leading in most animals to complete tumor regression (Figure 1E). The combination of both mAbs appeared to have no additional benefit (Figure 1E). The strong responsiveness to αPD-1 monotherapy clearly distinguishes MM008i from MM001i. Moreover, neither tumor line appeared to be immune-cold as evidenced by clear responses to αCTLA-4 monotherapy. As for MM001i tumors above, no therapeutic benefit was observed for MM008i in NSG mice in any immunotherapy treatment group, further consistent with an immune response-dependent mechanism of tumor regression (Figure 1F).

### Immunotherapy of MM001i and MM008i tumors generates immune memory

We next asked if immunotherapy treatment of MM001i and MM008i tumors triggers long-lasting immune memory. Mice were engrafted with MM001i and MM008i and, beginning day 7, treated as described above (workflows in Figure 2A,C). Mice demonstrating complete tumor regression following αPD-1 and/or αCTLA-4 treatment were allowed to rest for 90 days and then rechallenged through injection with the same tumor cell line at the same site as in the primary injection. Age-matched female WT mice, which had never been injected with tumor cells previously, were engrafted with MM001i and MM008i as controls. As expected, control mice with *de novo* injections developed mammary tumors in 21-28 days following MM001i or MM008i engraftment (Figure 2B,D). Conversely, all previously treated tumor-free mice rejected additional engraftment attempts, thus indicating that both MM001i and MM008i are capable of provoking long-term anti-tumor immune memory.

**Figure 2.**
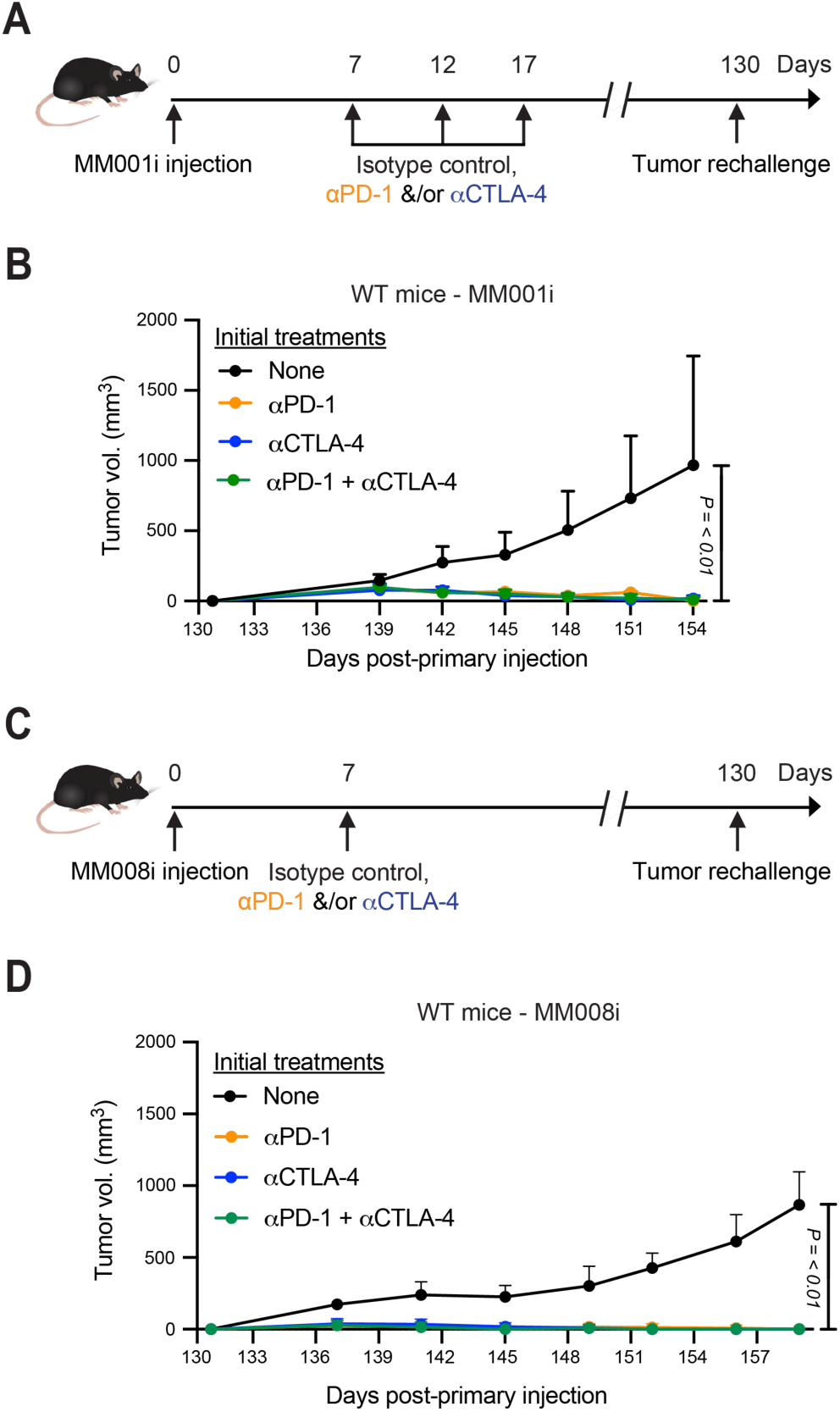
**MM001i and MM008i tumors elicit durable immune memory response post-immunotherapy treatments.** (**A**) Timeline of initial and subsequent MM001i tumor injections and treatment with αPD-1, αCTLA-4, or combination. (**B**) Tumor growth curves of MM001i following second tumor engraftment into WT female mice. n=4 animals per group (1 mouse for αPD-1) with two tumors per mouse; *P*-values using two-way ANOVA. (**C**) Timeline of initial and subsequent MM008i tumor injections and treatment with αPD-1, αCTLA-4, or combination. (**D**) Tumor growth curves of MM008i following second tumor engraftment into WT female mice. n=5 animals per group with two tumors per mouse; *P*-values using two-way ANOVA.

### CD3^+^ T cells are distributed throughout MM008i unlike MM001i tumors

Although MM001i and MM008i originated from the same genetic background, these two mammary tumor cell lines exhibit starkly different responses to αPD-1 treatment (Figure 1). To begin to investigate the underlying mechanism, immunohistochemistry was used to profile tumor-infiltrating immune populations including as B and T lymphocytes. Untreated MM001i and MM008i tumors were isolated after reaching the endpoint tumor volume of 1000 mm^3^ and stained for CD19 and CD3 expression. Based on CD19 staining, B cell infiltration was low in both MM001i and MM008i tumors (not shown). In contrast, both mammary tumors showed evidence for T cell infiltration with MM008i tumors exhibiting significantly more CD3^+^ T lymphocytes compared to MM001i tumors (Figure 3A,B; positive control splenic CD3^+^ cell staining in Supplementary Figure 3A). Moreover, T cells were localized throughout MM008i tumors versus predominantly around the peripheral margins of MM001i tumors (Figure 3A). Deeper T lymphocyte penetration is consistent with the possibility that this cell type may be responsible for the differential αPD-1 therapeutic outcomes of MM008i and MM001i tumors.

**Figure 3.**
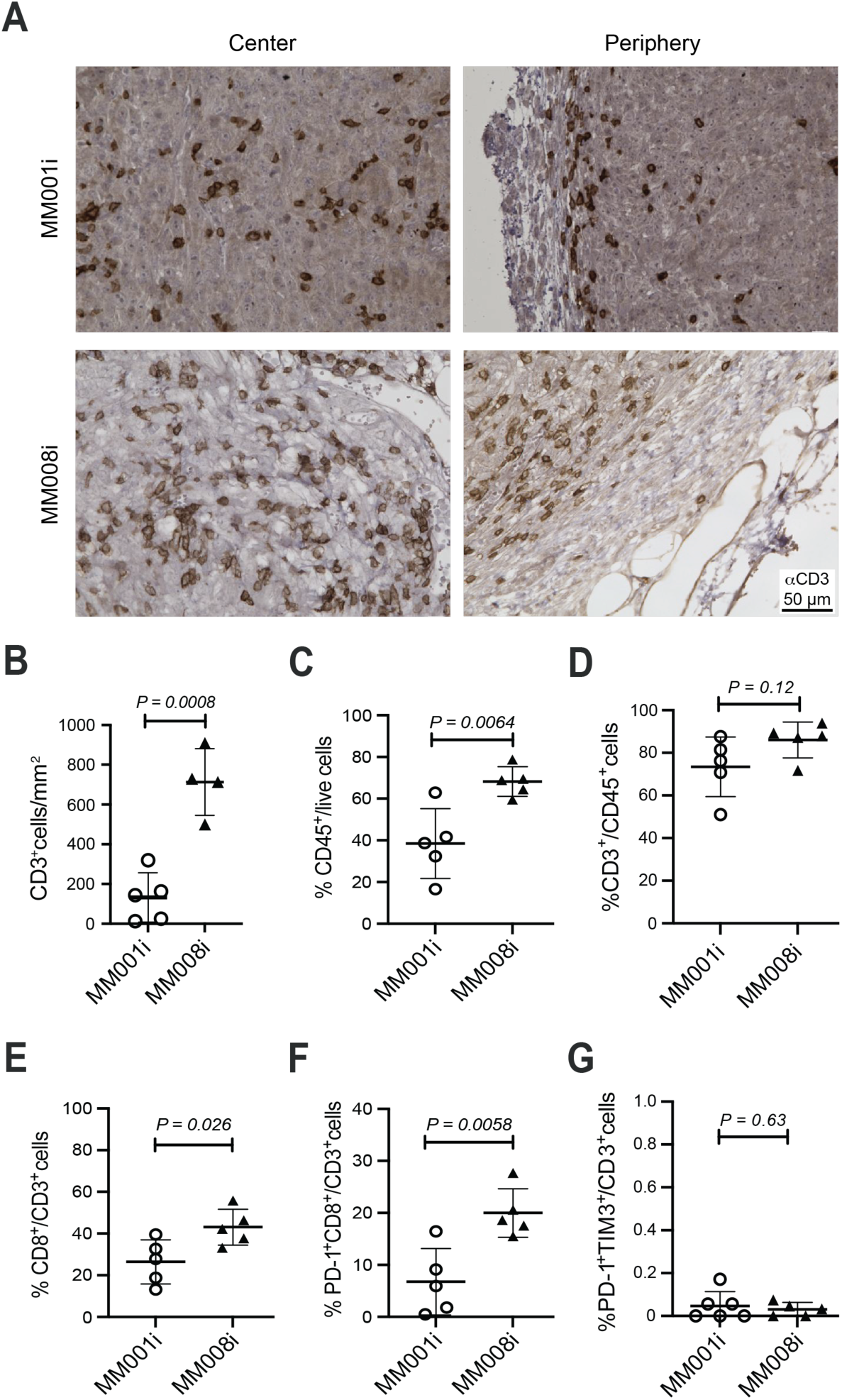
**MM008i tumors are enriched with total T cells and pre-exhausted PD-1^+^CD8^+^ T cells compared to MM001i tumors.** (**A**) Representative αCD3 IHC images of untreated MM001i and MM008i tumors. (**B**) Quantification of CD3^+^ T cells/mm^3^ in untreated MM001i and MM008i tumors using QuPath bioimage analysis software (5 random uniform regions of interest per tumor; n=5 tumors for MM001i, and 4 tumors for MM008i; *P*-value by unpaired Student’s *t-*test). (**C-G**) Immune cell infiltrates with the indicated cell surface markers as estimated by flow cytometry (n=4-5 tumors per group; *P*-value by unpaired Student’s *t*-test).

### Infiltrating CD8^+^ T cells in MM008i tumors

To further profile tumor-associated immune cells, flow cytometry was used to assess specific immune cell subpopulations in treatment-naïve primary tumors. Following live cell gating, a significantly higher percentage of CD45^+^ immune cells was identified in MM008i tumors versus MM001i tumors (Figure 3C; gating strategy and representative scatter plots in Supplementary Figure 4). There was also a trend toward higher CD3^+^ T cells out of CD45^+^ immune cells (Figure 3D) and, interestingly, a significantly higher percentage of CD8^+^CD4^-^ cytotoxic T cells in the CD3^+^ population of MM008i tumors (Figure 3E). Additionally, activated, pre-exhausted PD-1^+^TIM3^-^CD8^+^ T cells^12, 39^ were significantly higher, though no difference was observed for exhausted PD-1^+^TIM3^+^ T cells in MM008i compared to MM001i tumors (Figure 3F,G).^40, 41^ No significant difference was observed for CD11b^+^ monocytes, CD11b^+^Gr1^+^ myeloid-derived suppressor cells, or B220^+^ B cells (Supplementary Figure 3B,C,D). A trend towards a higher percentage of CD11b^+^CD11c^+^ dendritic cells and NK1.1^+^ natural killer cells was observed in MM001i tumors, though it was not statistically significant (Supplementary Figure 3E,F). These observations combine to indicate that the successful αPD-1 outcomes observed for MM008i tumor-engrafted animals may be mediated through pre-exhausted PD-1^+^TIM3^-^CD8^+^ T cells.

### Depletion of CD8^+^ cells abrogates αPD-1 treatment efficacy in MM008i

The aforedescribed results suggest that the αPD-1 responsiveness of MM008i may be due to infiltrating anti-tumor CD8^+^ T cells. To test this possibility directly, experiments were performed in which CD8^+^ T cells were immunodepleted from animals before and after MM008i tumor treatment with αPD-1 (workflow in Figure 4A). Tumors showed normal growth kinetics in WT animals treated with isotype control mAbs or an αCD8 mAb (Figure 4B). As above, αPD-1 treatment alone resulted in near-complete tumor regression (Figure 4B). However, three doses of αCD8 mAb (one before and two after αPD-1 treatment) were sufficient to blunt the therapeutic responsiveness of the αPD-1 therapy (Figure 4B). Immunohistochemistry confirmed depletion of CD8^+^ T lymphocytes from MM008i tumors (representative images in Figure 4C). These data combined to show that CD8^+^ T cells are an integral cell population required for MM008i tumor regression in response to αPD-1 therapy in WT mice.

**Figure 4.**
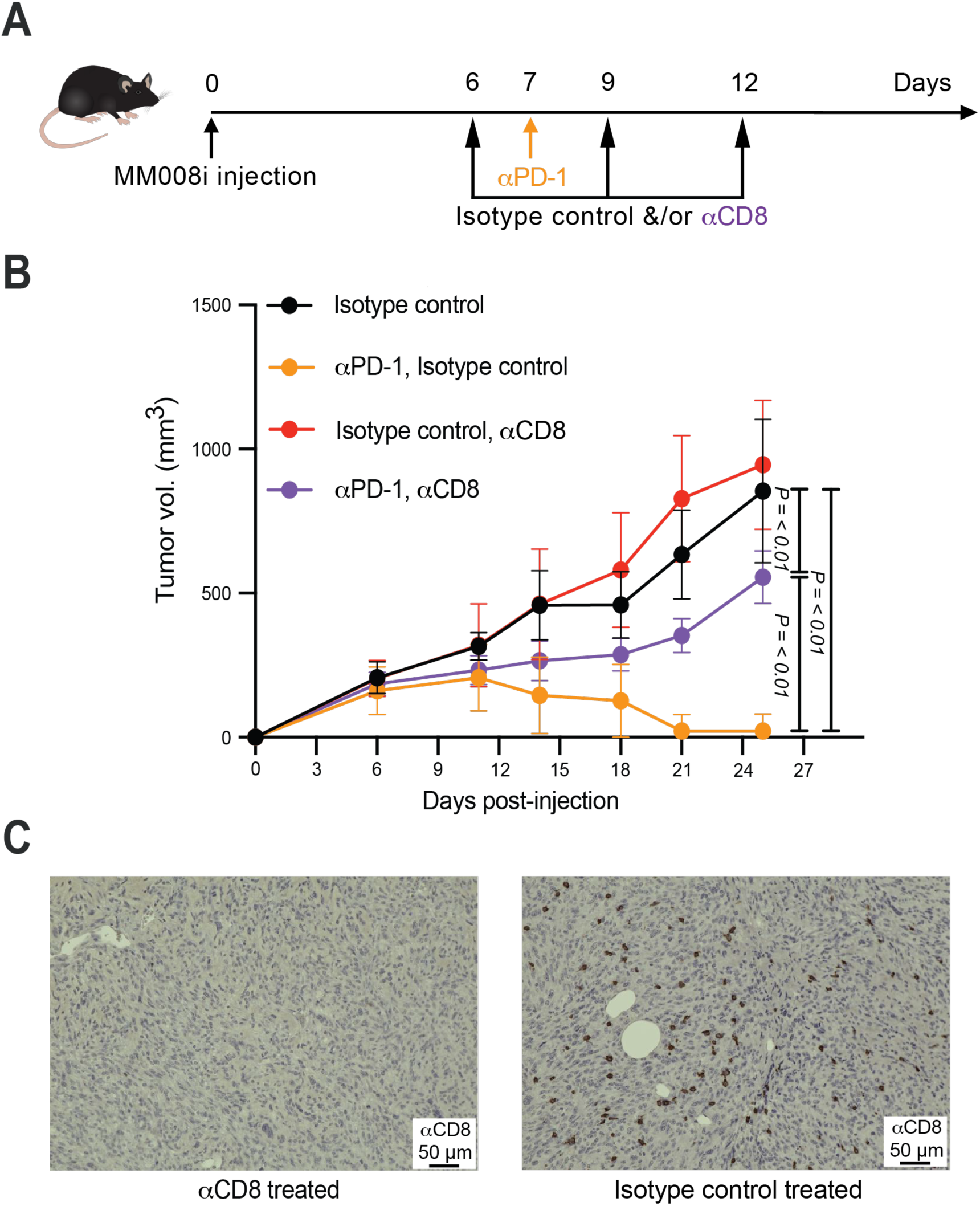
**αCD8 neutralizing antibody abrogates αPD-1 response in MM008i tumors.** (**A**) Timeline of MM008i tumor injections and treatments with αPD-1 ± αCD8 or isotype controls. (**B**) Tumor growth curves of MM008i following tumor injection in WT females in the indicated treatment groups. n=5 animals per group with two tumors/mouse; *P*-values using two-way ANOVA. (**C**) Reconstruction experiment showing representative αCD8 IHC images of MM008i tumors on day 13, immediately following treatment of animals with αCD8 neutralizing antibody or an isotype control antibody.

### RNA-sequencing of MM001i and MM008i cells

To identify tumor-intrinsic factors that contribute to the differential responsiveness to αPD-1, bulk RNA sequencing was done for cultured asynchronous MM001i and MM008i cell populations (2 independent cultures per line) and for untreated tumors from animals (3 independent tumors per line). A full list of expressed genes from cultured cells and untreated tumors is provided in Supplementary Table 1. First, as expected from initial genotyping above, no transcripts were detected for *p53* exons 2-10 confirming *p53* loss of function. Second, multiple breast cancer-associated genes were expressed at similar levels in both cell lines (*Brca1, Brca2, Mki67, Myc, Pik3ca*, *Pten*, *Rb1*) with no apparent disease mutations. Third, consistent with IHC results above, *Esr1* transcripts were not detected in either cell line. Fourth, in contrast to the IHC results above where HER2 was not detected, both lines appear to express similar levels of *Erbb2* RNA. This inconsistency may be due to an intrinsic cellular defect prior to translation as this protein was undetectable by IHC and unlikely to be retained intracellularly. Finally, modest levels of *Ar* and *Pgr* RNA were observed in MM001i cultured cells, and not in MM008i. Expression comparisons were less clear in bulk RNA from tumors due to a multitude of normal infiltrating cell types.

To evaluate broad expression differences between MM001i and MM008i cells and tumors, we used principal component analysis (PCA) to compare the overall transcription profiles for MM001i and MM008i in each growth condition (Supplementary Figure 5A). In general, cultured cells clustered closely together as anticipated. In comparison, tumor cells clustered far away from cultured cells and in more loosely associated groups consistent with differential immune responses. We next analyzed specific gene expression differences between MM001i and MM008i, identified based on a greater than 4-fold change and *P* values < 0.05 for cultured cells and < 0.01 for untreated tumors (Figure 5A; Supplementary Table 1). Among the highest differentially expressed genes are immune related genes, such as *Il11*, *Cxcl2*, *Il7*, *Il1rap*, *Ifnlr1*, and *Cxcl3* in tumors, *Tnf* and *Havcr2* (encoding TIM3) in cultured cells, and *Il1rl1*, *Nfgr*, and *Ptgs2* in both sets, which could potentially relate to the observed differences in response to αPD-1 therapy and in T lymphocyte infiltration.

**Figure 5.**
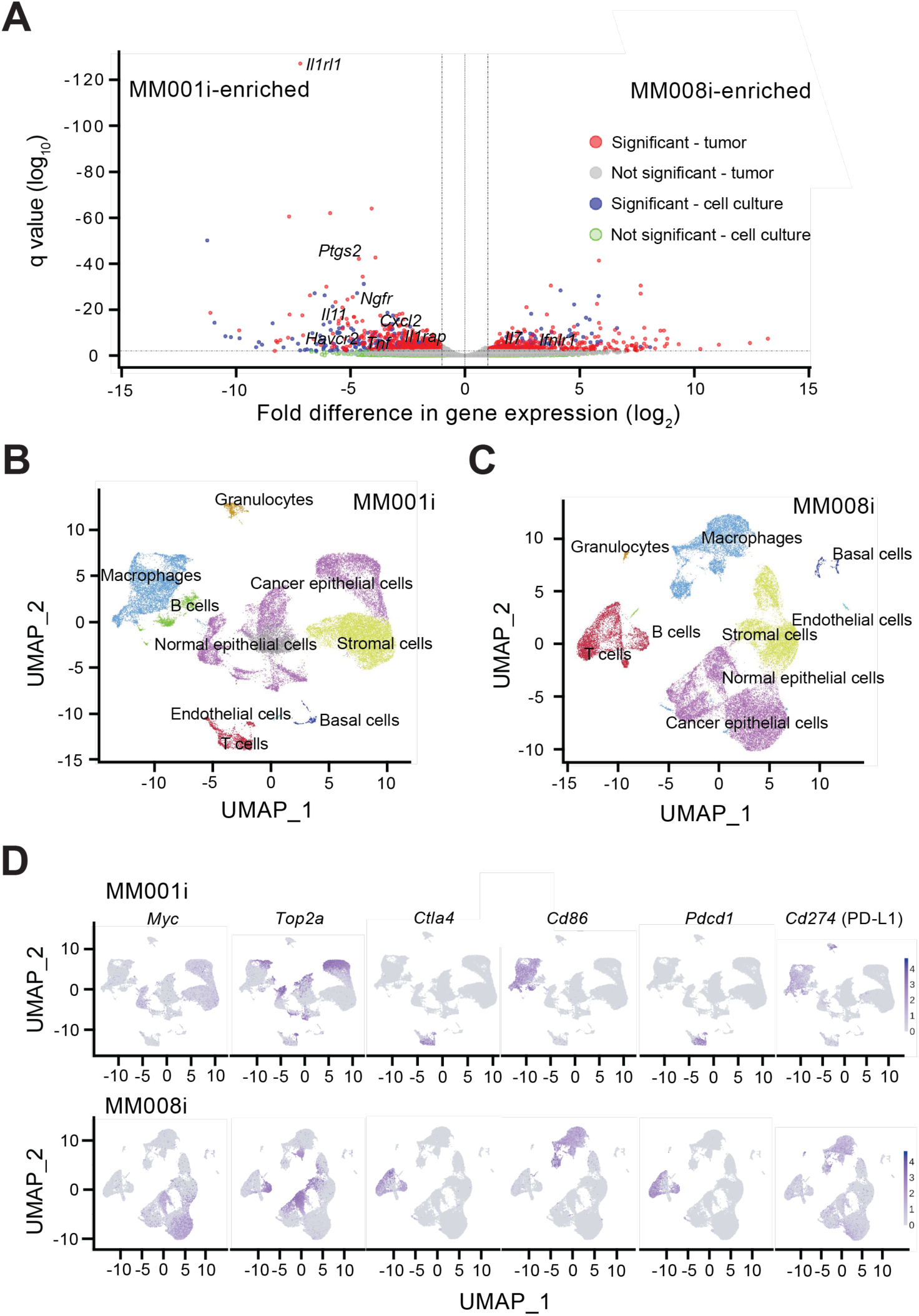
**Gene expression profiling of MM001i and MM008i tumors.** (**A**) Volcano plot depicting differentially expressed genes between MM001i and MM008i cells in culture and tumors *in vivo* by RNA-seq (filtered by log_2_-fold change > 2 and adjusted *P*-value <0.05 for cultured cells and < 0.01 for tumors). (**B-C**) UMAP clustering of MM001i and MM008i tumors, respectively, with cell type annotations by single-cell RNA-seq. (**D**) Expression distributions for *Myc*, *Top2a*, *Ctla4*, *Cd86*, *Pdcd1*, and *Cd274* at the single cell level in MM001i and MM008i tumors overlaid upon respective UMAPs.

To evaluate gene expression at the pathway level, we conducted KEGG signaling pathway analysis on the top 1000 expressed genes in MM001i and MM008i cells and tumors, focusing specifically on enriched pathways in immune signaling, cancer, and epithelial cell development (Supplementary Figure 5B; Supplementary Table 2). Overall, gene expression was very similar between MM001i and MM008i, particularly in epithelial cell signaling pathways. MM008i tumors were relatively enriched for immune related pathways such as antigen processing and presentation, and phagosome, with genes such as *H2-t23* and *B2m* being among the most strongly expressed. Conversely, MM001i tumors were relatively enriched for cancer related pathways such as extracellular matrix (ECM)-receptor interaction and PI3K-AKT signaling. These gene expression insights highlight the many similarities between MM001i and MM008i mammary tumor cell models and also point toward differences that may explain the differential responsiveness to immune therapy.

### Single-cell profiling of MM001i and MM008i tumors

Single cell (sc)RNA-seq analysis of untreated MM001i and MM008i tumors freshly isolated from mice revealed additional intrinsic and extrinsic features of each tumor cell line. Unsupervised clustering (UMAP) identified major cell types, including normal and cancer epithelial cells, endothelial cells, stromal cells, basal cells, and immune populations (Figure 5B,C). Epithelial cells in both MM001i and MM008i exhibited elevated expression of the oncogenes *Myc* and *Top2a*, indicating shared proliferative and malignant transcriptional programs (Figure 5D). *Ctla-4* (*CTLA-4*) and *Pdcd1* (*PD-1*) were highly expressed in T cells of both MM001i and MM008i, whereas the mRNA for the CTLA-4 ligand *Cd86* showed no significant upregulation in either MM001i or MM008i (Figure 5D). In contrast, *Cd274* (encoding PD-L1) was overexpressed specifically in epithelial cells of MM008i but not in epithelial cells of MM001i, implying specific immune evasion via the PD-1/PD-L1 axis in MM008i tumors (Figure 5D).

## Discussion

We report the development of two syngeneic murine mammary tumor cell lines that, despite originating from the same C57BL/6J genetic background, display differential *in vivo* responsiveness to αPD-1 checkpoint blockade. Specifically, the mammary tumor cell line MM008i tumors responds to αPD-1 therapy and MM001i does not. However, both lines are fully responsive to αCTLA-4 therapy, and tumor clearance associates with immunity to subsequent engraftment. Moreover, comparative immune cell profiling of engrafted tumors showed higher percentages of CD8^+^ cytotoxic T cells in MM008i tumors compared to MM001i tumors. These CD8^+^ T lymphocytes are required for response to αPD-1 in MM008i tumors, as depletion of this population abrogates therapy response. Another key difference we identified was significantly higher expression of PD-L1 on MM008i epithelial cells compared to MM001i. This strongly indicates that the PD-1/PD-L1 axis is a major mediator of immune evasion in MM008i, which is reflective of αPD-1 clinical observations in breast cancer where PD-L1 serves as a key biomarker for positive responses.^11, 42, 43^ However, despite these clear phenotypic differences between tumors, MM001i and MM008i are overall very similar at the gene expression level. We observed similar expression levels of commonly dysregulated breast cancer genes such as *Brca1, Brca2, Mki67, Myc, Pik3ca*, *Pten*, *Rb1*, and *Top2a*, and similarities in epithelial signaling pathways.

The mammary tumor models presented help fill an important gap in syngeneic systems for preclinical immune therapy testing and mechanistic studies. Syngeneic mammary tumor models are rare, with only a few described to date.^14, 18, 21, 24–26, 44^ E0771, which has been around for 70 years and with unknown passage number, has conflicting reports on ER hormone receptor status.^18^ Other lines, such as PyMT,^24^ AT-3,^25^ and WT145^26^ rely on either transgene insertion or chemical carcinogens to drive cancer formation, processes that do not reflect cancer development in human patients. Further, it would be difficult to compare results across lines given substantial differences between each in genetic background, growth kinetics, and immune phenotype, limiting the potential for comparative analysis. To address some of these limitations, recent efforts have focused on developing new models that better recapitulate the immunological characteristics of human disease.^45^ However, many of these models are poorly responsive to immunotherapy, owing to low tumor mutational burdens, unless further modified by induced mutagenesis or *Brca1* deficiency.^45^ Our new models, which exhibit clear responses to αCTLA-4 and differential responses to αPD-1, provide a unique platform to dissect mechanisms of immunotherapy resistance and test additional therapies (alone and in combination).

In particular, we envision our mammary tumor models being used to test other therapies in combination with αPD-1 for potential dosage de-escalation of MM008i and to potentially trigger αPD-1 responsiveness to MM001i. Our TNBC models also may be used to dissect the tumor microenvironment itself, particularly focusing on extracellular matrix signaling and immune infiltration. Here, we demonstrate considerable differences in T cell infiltration between our MM001i and MM008i tumors. However, we have not directly investigated the specific mechanisms which permit greater immune infiltration in MM008i tumors than MM001i tumors. We anticipate future studies looking more deeply into these two cell lines immunologically, particularly the factors that influence whether a tumor is “immune hot” or “immune cold”. By identifying some of these molecular determinants, we may be able to identify combination therapy approaches to make previously resistant tumors sensitive to immunotherapies.

## Acknowledgments

We thank Francesca Manara (Memorial Sloan Kettering Cancer Center), Josephine Taverna (UT Health San Antonio), and Ratna Vadlamudi (UT Health San Antonio) for thoughtful feedback, and Emily Law, Jordan Naumann, and Angel Dominguez for early contributions to this project. We are also grateful to Zhao Lai at the UT Health San Antonio Genome Sequencing Facility for assistance with library preparation and sequencing and to the South Texas Research Histology Laboratory for help with mouse specimen processing.

## Author contributions

Reuben S. Harris, Harshita B. Gupta: Conceptualized the project; Thomas J. Kalantzakos, Harshita B. Gupta, Cameron Durfee, Anusha Soni, Benjamin Troness: Conducted *in vivo* experimentation; Xingyu Liu, Joshua Proehl, Harshita B. Gupta, Ian Tamayo: Performed immunohistochemistry; Yufan Zhou, Xingyu Liu, Nuri Alpay Temiz: Contributed to computational analyses; Thomas J. Kalantzakos, Harshita B. Gupta: Performed cell-based experiments; Thomas J. Kalantzakos, Harshita B. Gupta, Reuben S. Harris: Drafted the manuscript with all authors contributing to manuscript proofing and revision.

## Data availability statement

The murine mammary tumor cell lines generated in this study, MM001i and MM008i, are available through the American Type Culture Collection or written request. Primary bulk and single-cell RNAseq data sets are available through GEO (GSE305800 and GSE305842; respectively https://www.ncbi.nlm.nih.gov/geo/query/acc.cgi?acc=GSE305800 and https://www.ncbi.nlm.nih.gov/geo/query/acc.cgi?acc=GSE305842. Supplementary Tables 1 and 2 are available on Figshare (doi: 10.6084/m9.figshare.30468098) using the link https://figshare.com/s/cd92140dd5a6f2f5a9d4. All other primary data are presented in the manuscript. A preprint of this work is also available on *bioRxiv* (https://www.biorxiv.org/content/10.1101/2025.09.18.677171v1).

