## Supplementary Figures 1-5 for "Murine models for triple-negative breast cancer with differential responsiveness to immunotherapy"

**Supplementary Table 1.** Gene expression of MM001i and MM008i cultured cells and tumors. See separate xlsx file.

**Supplementary Table 2.** Top 1000 expressed genes in bulk RNA-seq pathway analyses for MM001i and MM008i cultured cells and tumors. See separate xlsx file.

**Supplementary Figures 1-5.** See following pages.

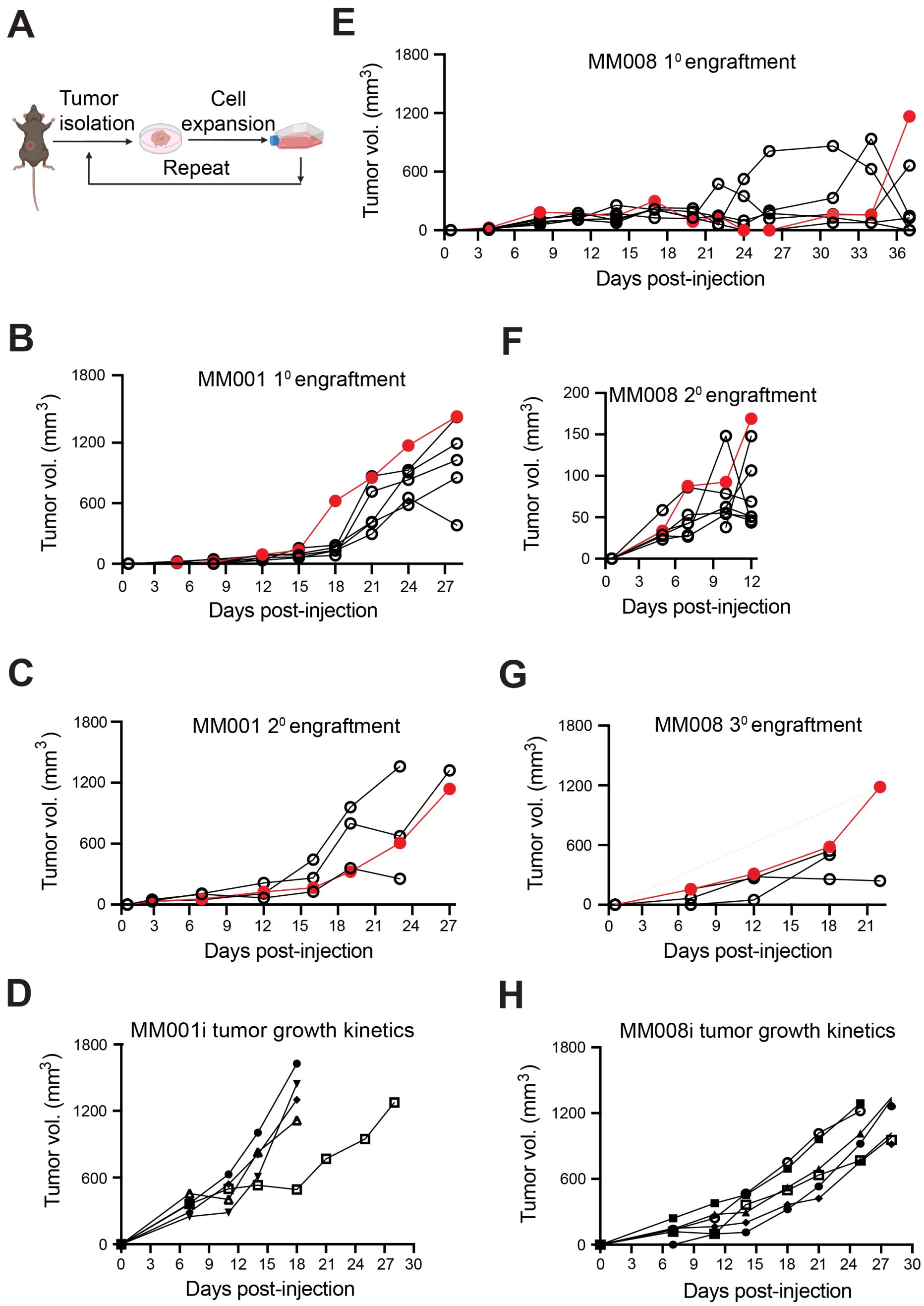


**Supplementary Figure 1.** Development of MM001i and MM008i cell lines by serial passage in WT females.

(**A**) Workflow schematic of serial passage, where tumors are isolated, grown *in vitro*, and then re-engrafted in into naïve WT mice (created with BioRender.com).

(**B-D**) Primary, secondary, and tertiary (final) tumor growth curves for MM001, ultimately yielding MM001i (tumor selected for expansion in red).

(**E-H**) Primary, secondary, and tertiary, and quaternary (final) tumor growth curves for MM008, ultimately yielding MM008i (tumor selected for expansion in red).

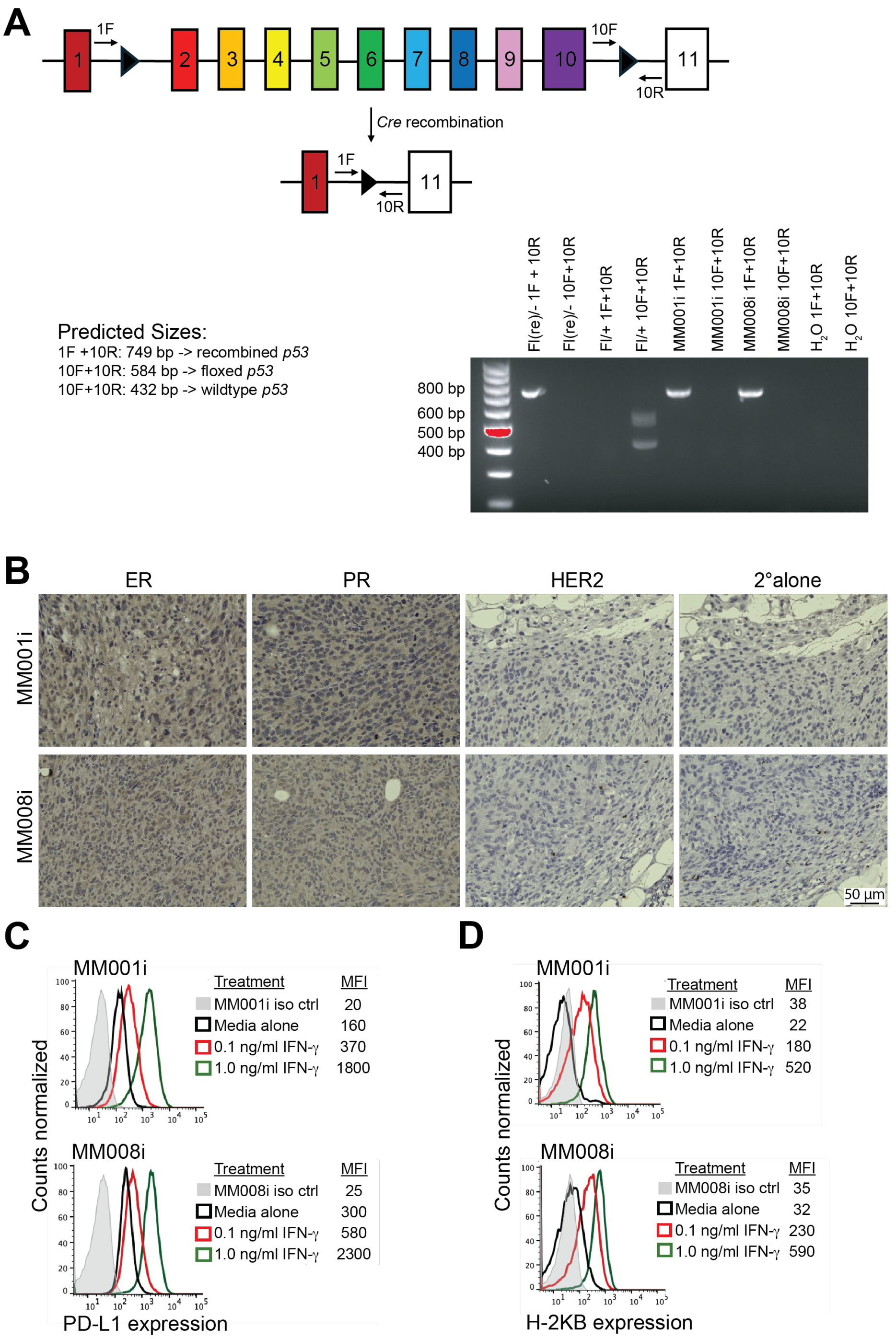


**Supplementary Figure 2.** Characterization of MM001i and MM008i cells.

(**A**) PCR strategy to assess the *p53* genotype of MM001i and MM008i cells.

(**B**) Representative IHC of hormone receptors ER, PR, and HER2 in MM001i and MM008i tumors.

(**C-D**) Flow cytometry expression levels of PD-L1 and H-2KB in the absence and presence of stimulation by IFN-γ.

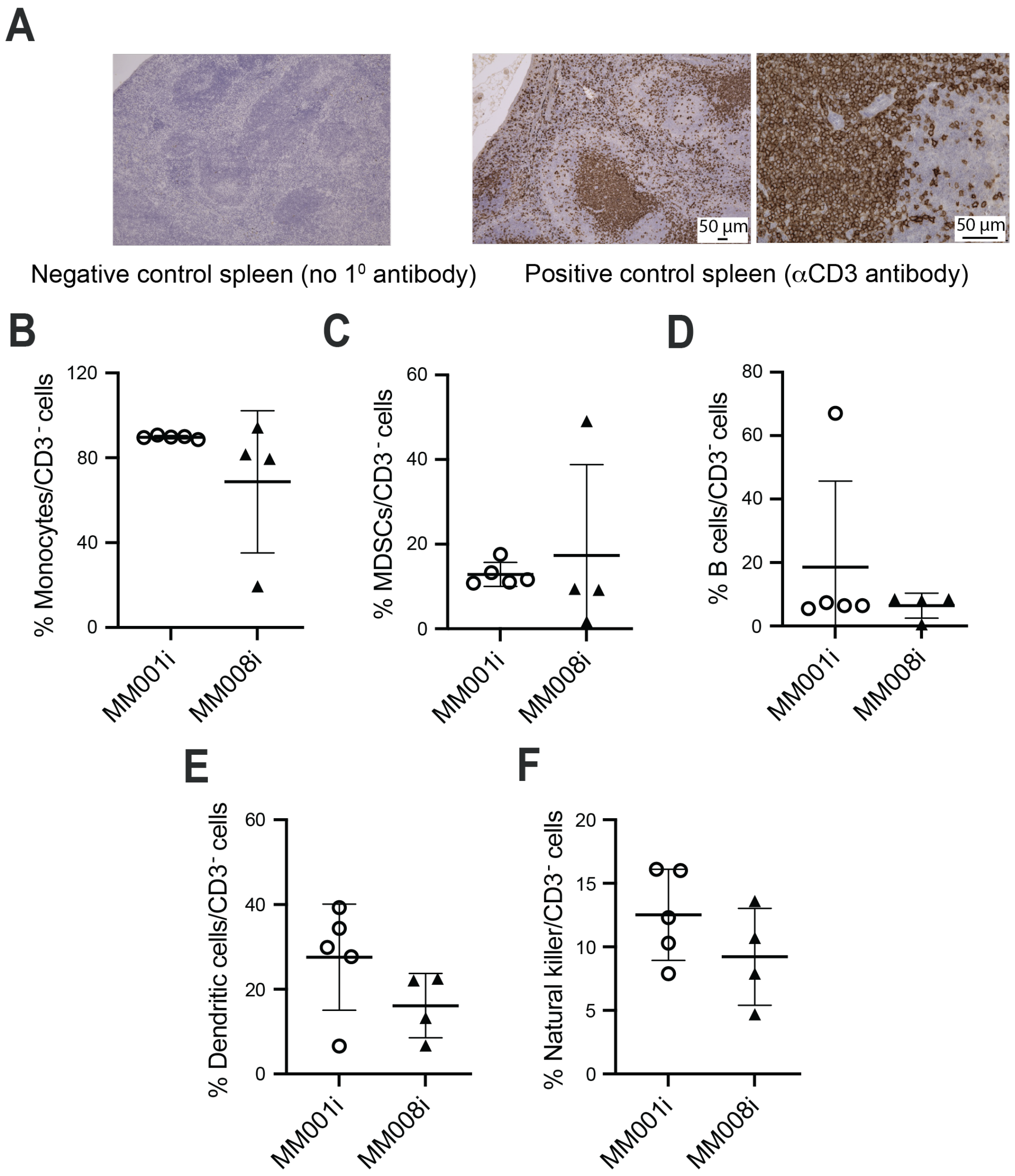


**Supplementary Figure 3.** Supplementary immune profiling of MM001i and MM008i tumors. (**A**) Negative (no primary antibody) and positive control ($\alpha$CD3) staining of splenic tissue. (**B-F**) Percent monocytes, myeloid-derived suppressor cells (MDSCs), B cells, dendritic cells, and natural killer cells relative to $CD3^-$ non-T cell immune population.

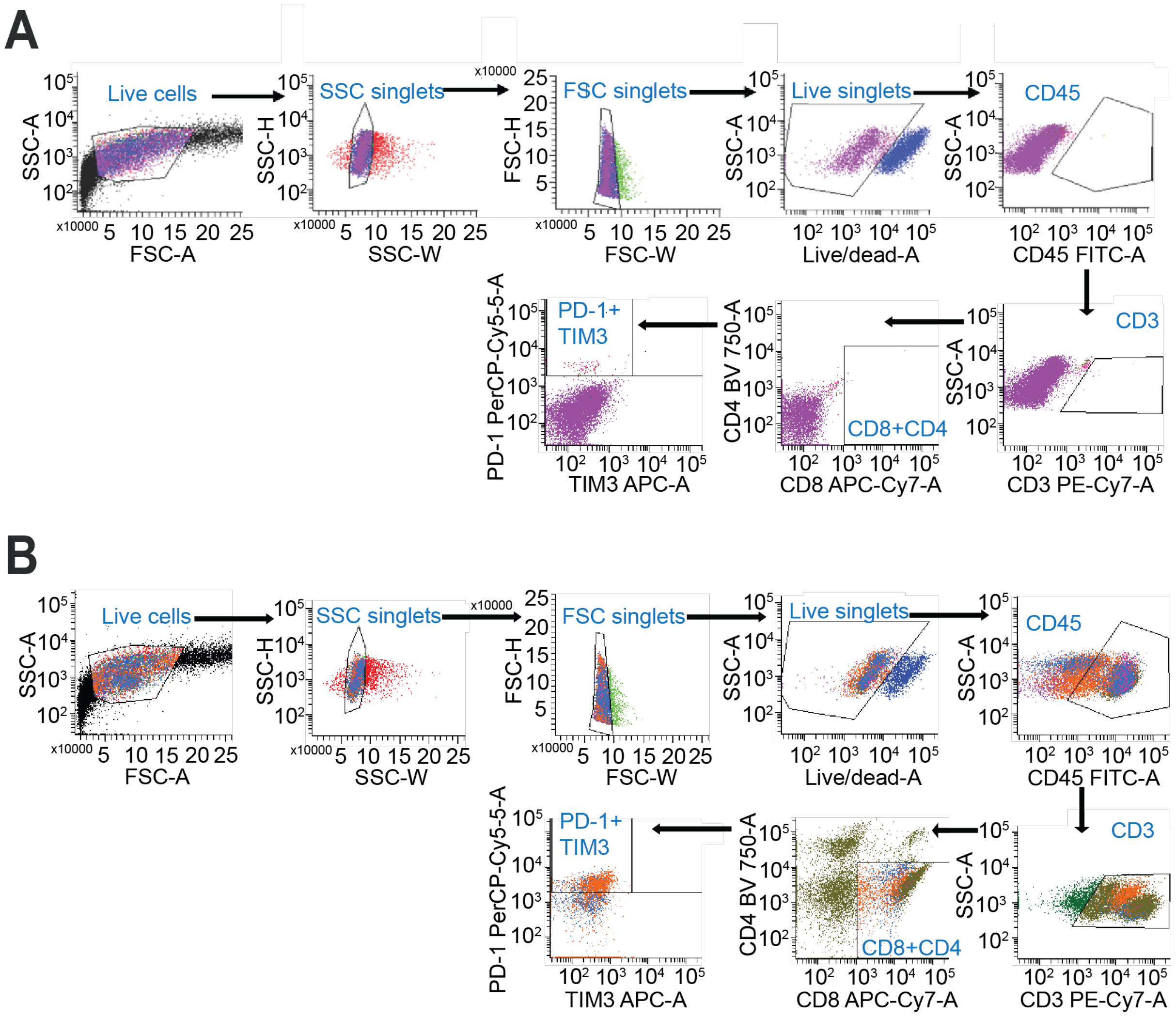


**Supplementary Figure 4.** Representative gating and flow cytometry scatter plots for analyzing immune infiltration in MM001i and MM008i tumors

(**A**) Representative gating strategy with MM008i stained with isotype control antibodies to establish positivity thresholds for identification of pre-exhausted PD-1$^+$TIM3$^-$ CD8$^+$ T cells. Gating was performed sequentially on singlets (FSC and SSC), live singlets (Zombie live/dead staining), leukocytes (CD45$^+$), T cells (CD3$^+$), CD8 T cell subsets (CD8$^+$CD4$^-$), and pre-exhausted CD8 T cells (PD-1$^+$TIM3$^-$).

(**B**) Representative flow cytometry scatter plots to identify T cell subpopulations in an MM008i tumor, resulting in identification of pre-exhausted PD-1$^+$TIM3$^-$ CD8$^+$ T cells.

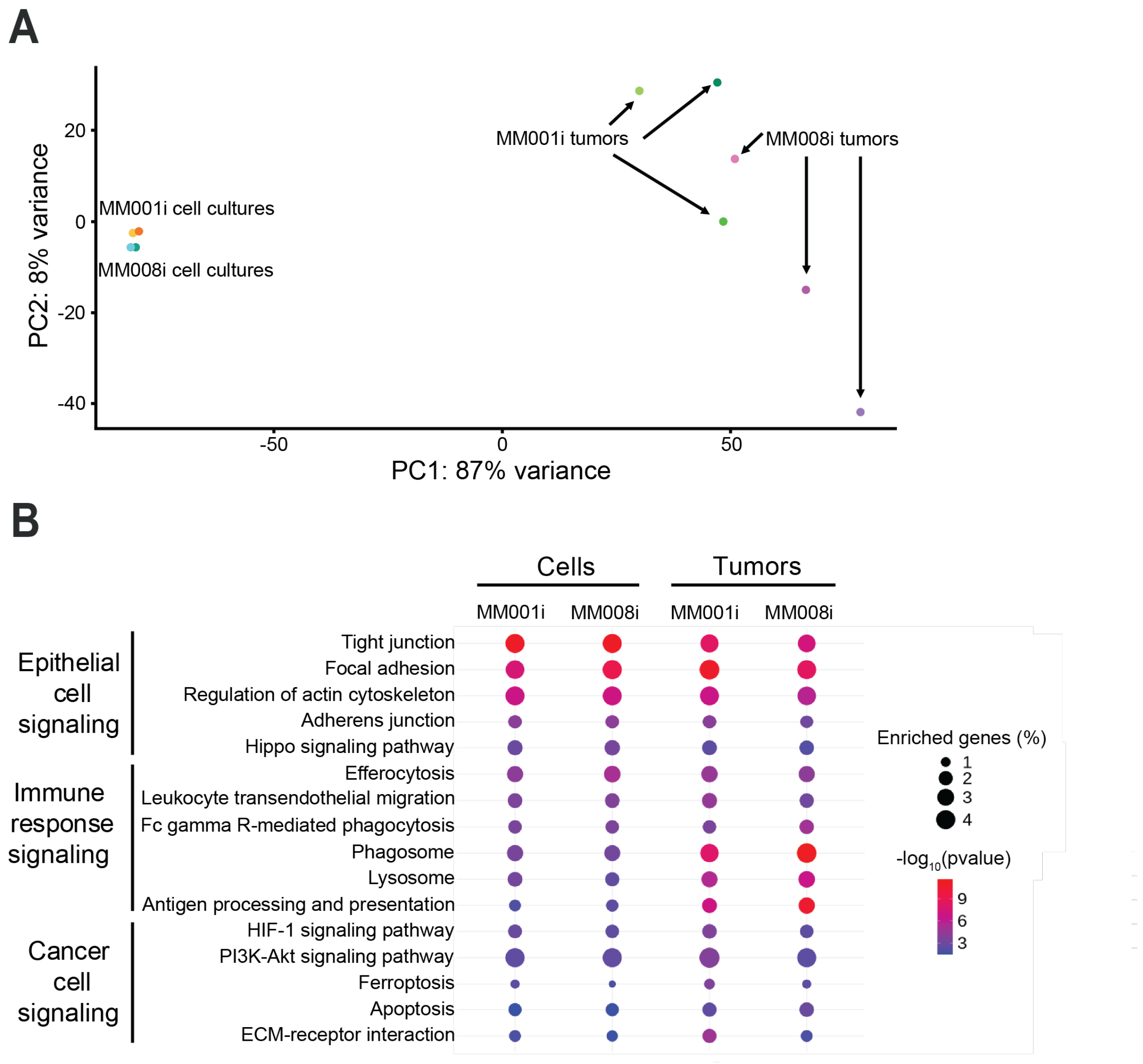


**Supplementary Figure 5.** Analysis of MM001i and MM008i cultured cells and tumors *in vivo* by RNA-seq.
(**A**) Principal component analysis comparing bulk RNA-seq expression profiles in MM001i and MM008i cells and tumors.
(**B**) Pathway enrichment in the top 1000 expressed genes by bulk RNA-seq in MM001i and MM008i cultured cells and tumors by KEGG signaling enrichment analysis, grouped by pathway type.
